# The Protein Common Assembly Database (ProtCAD) – A comprehensive structural resource of protein complexes

**DOI:** 10.1101/2022.08.15.503938

**Authors:** Qifang Xu, Roland L. Dunbrack

## Abstract

Proteins often act through oligomeric interactions with other proteins. X-ray crystallography and cryo-electron microscopy provide detailed information on the structures of biological assemblies, defined as the most likely biologically relevant structures derived from experimental data. In crystal structures, the most relevant assembly may be ambiguously determined, since multiple assemblies observed in the crystal lattice may be plausible. It is estimated that 10-15% of PDB entries may have incorrect or ambiguous assembly annotations. Accurate assemblies are required for understanding functional data and training of deep learning methods for predicting assembly structures. As with any other kind of biological data, replication via multiple independent experiments provides important validation for the determination of biological assembly structures. Here we present the Protein Common Assembly Database (ProtCAD), which presents clusters of protein assembly structures observed in independent structure determinations of homologous proteins in the Protein Data Bank (PDB). ProtCAD is searchable by PDB entry, UniProt identifiers, or Pfam domain designations and provides downloads of coordinate files, PyMol scripts, and publicly available assembly annotations for each cluster of assemblies. About 60% of PDB entries contain assemblies in clusters of at least 2 independent experiments. All clusters and coordinates are available on ProtCAD web site (http://dunbrack2.fccc.edu/protcad).

## INTRODUCTION

Many proteins function as homo- and heterooligomers by interacting with one or more copies of themselves or other proteins. The three-dimensional structures of these oligomers are referred to as biological assemblies (1). The biological assembly of an entry in the Protein Data Bank (PDB) (2) is often defined as the largest functional form present in an experimental structure obtained by X-ray crystallography, nuclear magnetic resonance (NMR), cryo-electron microscopy (EM), or other experimental methods. EM and NMR experimental data provide direct information on complexes or contacts needed to build an assembly. For crystal structures (~87% of the current PDB), the biological assembly is often ambiguous since the PDB provides several biological assemblies which are defined either by authors of a structure or predicted by the program Protein Interfaces, Surfaces and Assemblies (PISA) (3). The error rate in biological assembly annotations in the PDB has been estimated from 6 to 14% (4,5). The Protein Data Bank in Europe (PDBe) lists one PDB biological assembly as the “preferred” biological assembly primarily based on QSbio (6).

Several computational methods have been developed to identify the correct biological assembly within crystals. The PISA method, developed in 2007, has been a very successful program to predict biological assemblies from crystals (3,7). It emphasizes the necessity of considering the thermodynamics of the formation of entire assemblies and an evaluation of the chemical thermodynamics of assembly formation including entropy and desolvation. EPPIC3, developed in 2018 by Bliven et al., rigorously implements rules for the possible assemblies that can be found in a crystal – building neighboring unit cells, closed assemblies with point group symmetry, and full coverage of the crystal with a single type of assembly (8). It uses evolutionary conservation in the interfaces within each assembly to predict which assembly is most likely to be biological. QSbio determines biological assemblies in the PDB from pairwise structural alignments of homologous assemblies limited to the PDB’s deposited assemblies and predictions from PISA and EPPIC (6). Korkmaz et al. developed a procedure to correct biological assemblies in the PDB by comparing the symmetry and stoichiometry of an assembly with evidence from author annotations, text mining of the literature, and predictions from EPPIC and PISA across homologous PDB entries (9). They concluded that 83% of PDB entries may be considered to have the “correct” assembly annotation, while the remaining 17% can be split into 9% wrong and 8% inconclusive. Machine learning techniques have also been utilized in biological assembly prediction. Inference of Protein Assembly in Crystals (IPAC) utilized a set of 10 features of interfaces – based on packing, surface complementarity, surface area, and solvation energies – with a naïve Bayes classifier model to predict full quaternary structures in crystal structures (10). They focused particularly on the prediction of weak, transient oligomeric structures, which are poorly predicted by most methods. Prediction of protein assembly structures has become an important part of the CASP protein structure prediction experiments (11,12). Numerous methods have been developed for the simpler problem of identifying biological interfaces in crystals instead of full assemblies (13–19).

A data-driven approach to this problem is the observation that biologically relevant assemblies are often observed across crystal forms of the same or homologous proteins (6,20,21). For example, the enzyme, porphobilinogen synthase (PBGS), forms octamers in 8 different crystal forms for 8 different UniProt sequences (22). The RCSB PDB provides a very useful service that finds similar assemblies for a given query in the deposited biological assembly structures present in the PDB (23). In the Protein Common Interface Database (ProtCID), we have previously applied this principle by clustering both homodimeric and heterodimeric protein-protein (24) and domain-domain (25) interfaces observed in crystal structures of homologous proteins. In ProtCID, to provide evidence of the biological relevance of interfaces, we counted crystal forms (CFs, crystals with homologous proteins but different unit cells and/or space groups) rather than just entries. This is because some proteins are crystallized over and over in the same crystal form, which does not provide independent data on interfaces or assemblies. We have shown previously that the probability of biological relevance of an interface is correlated with the number of crystal forms in the PDB that contain the interface (20). This is a principle commonly used by crystallographers in determining the most likely assembly in their structures.

While data on protein interfaces across the PDB is useful, many such interfaces are clearly part of larger assemblies. Homooligomeric protein assemblies usually have cyclic symmetry or dihedral symmetry, and the interfaces may be either asymmetric (heterologous) or symmetric (isologous) (26). *C2* homodimers contain one isologous interface, while larger cycles (*Cn*) contain *n* heterologous interfaces that produce the cycle. Dihedral symmetry structures of type *Dn* are built from isologous interactions (and interfaces) of two *Cn* cycles. This leads to a simple way to determine whether two assemblies are the same or different. For example, a pair of homologous *D3* (dihedral) hexamer structures must have at least two common interfaces, one of which is heterologous (around the cycles) and one of which is isologous (between one cycle and the other). Some assemblies may have forms of symmetry such as tetrahedral, octahedral, or dodecahedral. Many assemblies are asymmetric. This includes complexes of single copies of non-homologous proteins but also some homooligomers such as the activating homodimer of the EGFR kinase domain (27) and filamentous proteins such as actin.

In this paper, we present the Protein Common Assembly Database (ProtCAD), which contains clusters of assemblies of homologous proteins observed in multiple independent experiments, such as unique crystal forms and individual cryo-EM and NMR experiments. We use the term “Crystal forms” (CFs) in this paper to refer to independent experiments that provide evidence in favor of the structure of biological assemblies. CFs include independent cryo-EM and NMR experiments and clusters of crystallographic structures of the same proteins in the same unit cell. This is consistent with our previous nomenclature (20,24,25), since the majority of structures are still crystallographic. We build interactions within protein crystals using crystallographic symmetry operators and identify all unique viable assemblies that can constitute the entire crystal using EPPIC. In this way we are not reliant on the assemblies deposited by authors in the PDB, although we do include all of the PDB-annotated assemblies in the data set for clustering. We first group the assemblies across crystal forms and EM/NMR structures of homologous proteins (or protein complexes) by stoichiometry and symmetry (e.g., we only compare C3 to C3 assemblies and can skip comparing C3 to C4, etc.), and then check whether the necessary isologous and heterologous interfaces are shared We check the interfaces from largest to smallest. This is a very fast and accurate way of comparing and clustering assemblies, since it requires only lists of homologous sequence positions which we derive by aligning sequences to Pfam HMMs and pairwise interface similarities (already available in ProtCID). It does not require structure alignment of large assemblies.

Assemblies occurring in two or more crystal forms (including EM and NMR experiments) comprise a common assembly cluster in ProtCAD. About 100,000 entries (59% of all PDB entries) appear in clusters with at least two crystal forms. About 65,000 are contained in clusters of at least five crystal forms and the PDB annotations have the same assembly in about 85% of these. In addition, the observation of many crystal forms without a common assembly (dimer or larger) is good evidence in favor of a monomeric protein. We annotate whether the PDB, EPPIC, and PISA have the clustered assembly, which may suggest that the assembly is biologically relevant (by the criteria of author annotations, sequence conservation in interfaces, or biophysical properties, respectively). For each UniProt sequence, we also determine what percentage of the available crystal forms of that protein contain the same assembly, an additional indication of the biological relevance of the assembly. With a click of a single button, the user can download all the structures of a particular assembly across PDB entries and PyMOL scripts for aligning and visualizing them. All clusters and coordinates are available on http://dunbrack2.fccc.edu/protcad.

## MATERIAL AND METHODS

### Assembly data set

EPPIC assembly files in json format were downloaded from the EPPIC web server (https://www.eppic-web.org/) by its REST API (rest/api/v3/job/assemblies/{PDB ID}). EPPIC data were parsed and inserted into a Firebird database (https://firebirdsql.org). EPPIC assemblies were generated from the chain components and symmetry operators defined in the json files and the coordinates in PDB mmCIF files. We excluded those EPPIC assemblies if any components of an assembly are not connected, which occurs in at least one assembly for 24% of PDB entries. PDB biological assemblies were added into the assembly pool if they were not present in one of the EPPIC assemblies.

### Pfam architectures

Pfam assignments for all chains in the PDB were obtained from our PDBfam database (28). A chain of an assembly is represented by a “chain Pfam architecture,” such as “(Cyclin_N)_(Cyclin_C),” with each Pfam in parentheses and connected by underscores. An “entry Pfam architecture” is simply the concatenation of the Pfam architectures of the unique Pfam architectures in the entry, e.g., (Pkinase)(Cyclin_N)_(Cyclin_C). A “group” of entries or crystal forms is defined by those with the same entry Pfam architecture. An “assembly Pfam architecture” is composed of the unique chain Pfam architectures in the entry with copy numbers (e.g., “(Pkinase)2”). A Pfam architecture for an assembly can contain chains with a single Pfam architecture (same-Pfam arch) or with two or more different Pfam architectures (diff-Pfam arch). A same-Pfam architecture group can contain hetero-oligomeric assemblies. For instance, the *C1* hetero-tetramer (*C1-A2BC*) of PDB: 7CE3 is composed of four chains of three UniProts (IDH3A_HUMAN (two copies), IDH3B_HUMAN and IDH3G_HUMAN). All chains have one Pfam: Iso_dh domain. The assembly belongs to (Iso_dh) Pfam architecture and has *A4* stoichiometry in terms of the (Iso_dh)4 Pfam architecture.

### Connecting interfaces of an assembly

All chain-chain interfaces were identified in each assembly as chain pairs with at least ten pairs of C_β_ atoms with distance ≤ 12 Å and at least one atomic contact within 5 Å, or at least five atomic contacts ≤ 5 Å. The similarity of interface pairs was calculated by the Q score described by Xu *et al*. (20), which is equal to a distance-weighted count of the common contacting residue pairs in two interfaces divided by the total number of unique residue pairs within the two interfaces (similar to a Jaccard index (29)). For each assembly, we selected a list of unique connecting interfaces to represent the assembly. For instance, for a *Cn (n* ≥ 2) homo-oligomeric assembly, there is only one type of connecting interface that repeats around the cycle. For an ideal *D2* homo-tetrameric assembly, there are two unique connecting interfaces, one within a symmetric homodimer and one between the homodimers. The procedure to identify the unique connecting interfaces of an assembly is described as follows.

For each assembly:

~~~
1. Calculate all pairwise chain-chain interfaces within the assembly
2. Calculate similarity (Q score) of all pairs of interfaces
3. Cluster interfaces by pair-wise Q score ≥ 0.75
4. Select the interface with the smallest ID in each interface cluster to define the set *{unique interfaces}*
5. Sort *{unique interfaces}* in descending order of surface area values
6. For each interface cluster in *{unique interfaces}*:
       Add representative interface to *{unique connecting interfaces}*
       For each member of cluster:
              If *{connected chains}* is empty:
                       Add two chains of interface to *{connected chains}*
              Else:
                       If one of the chains is already in *{connected chains}*: add the other
                       to *{connected chains}*
       If all chains are in *{connected chains}*: Stop
7. Return the sets *{unique interfaces}* and *{unique connecting interfaces}*.
~~~

The If-Then-Else statement ensures that all the chains are connected, since after the first two chains, chains can only be added if they are connected to a chain already in the set *{connected chains}*.

### Binary similarity and clustering of assemblies

The symmetry of each assembly is obtained from EPPIC but is updated with the AnAnaS program (30) for hetero-multimers with different proteins but the same Pfam architectures. In this study, the stoichiometry of these hetero-multimers is *An*, and belongs to the same assembly Pfam architecture as *An* homo-multimers of the same Pfam. The AnAnaS program gives the global symmetry (or pseudo-symmetry) for a hetero-multimer, while EPPIC determines the symmetry based on the unique sequences in each assembly. For instance, the hexameric assemblies (*A6*) of the Pfam architecture (AA_kinase) contain 47 homo-hexamers (*A6* with same UniProt) and 21 hetero-hexamers (*A3B3* with two UniProts: MOSA_AZOVD and MOSB_AZOVD). EPPIC has C3 for these 21 hetero-hexamers and *D3* for the 47 homo-hexamers, while AnAnaS has *D3* symmetry for all of the hetero-hexamers and homo-hexamers.

The similarity of two assemblies is binary: either they are the same (1) or different (0). The algorithm for a given assembly Pfam architecture and stoichiometry is as follows:

~~~
1. Calculate similarity scores between the *{unique interfaces}* of two assemblies
2. Determine whether the *{unique connecting interfaces}* of assembly 1 are all present
in the *{unique interfaces}* of assembly 2. If Yes, S12=1; else S12=0.
3. Determine whether the *{unique connecting interfaces}* of assembly 2 are all present
in the {*unique interfaces*} of assembly 1. If Yes, S21=1; else S21=0.
4. The assemblies are similar if S = S12 x S21 = 1.
~~~

Single linkage hierarchical clustering was used to cluster assemblies. Assemblies with the same assembly Pfam architecture (including stoichiometry) and symmetry with mutual binary similarity = 1 are placed into one cluster.

### Phylogenetic trees

Phylogenetic trees are built by ArboDraw (https://dunbrack.fccc.edu/retro/ArboDraw/). The evolutionary distances of UniProt sequences were generated by the Clustal Omega web server https://www.ebi.ac.uk/Tools/msa/clustalo/ with default settings.

## RESULTS

### Protein common assembly database (ProtCAD)

As of March 28, 2022, there are 10,677 Pfam families (v34) represented in protein sequences in the PDB. Chains in the PDB can be designated by their chain Pfam architecture, such as (Cyclin_N)_(Cyclin_C). Entries may be grouped by the “entry Pfam architecture,” which designates the unique Pfam architectures of the sequences in the entry, e.g. (Cyclin_N)_(Cyclin_C)(Pkinase). Note that different protein sequences in a single structure may have the same chain Pfam architecture. An “assembly Pfam architecture” includes copy numbers, when the copy number is greater than 2, e.g. (Cyclin_N)_(Cyclin_C)2(Pkinase)2. We refer to assemblies with the same entry Pfam architecture, but potentially different stoichiometries, as a “group”. A group can consist of a single Pfam architecture (“same-PfamArch”) or more than one distinct Pfam architectures (“diff-PfamArch”). There are 14,954 same-Pfam architectures and 16,766 diff-Pfam architectures for a total of 31,720 Pfam architectures which involve 179,228 PDB entries and 57,032 proteins (UniProts) in the current database.

We used EPPIC (8) to derive assemblies for each structure in the PDB. EPPIC defines valid assemblies for crystals as single assemblies that cover the whole crystal and are “closed” (i.e., excluding filamentous assemblies). We obtained 1,452,301 assemblies for 185,085 PDB entries from EPPIC (161,724 crystal structures; 9,394 cryo-EM structures, and the rest NMR or other experimental methods). Unfortunately, 51.4% of EPPIC assemblies are not fully connected, with one or more chains not having any contacts with other chains in multi-chain assemblies. This phenomenon occurs in 69.1% of EM entries and 22.8% of crystallographic entries. About a quarter of the disconnected assemblies are due to peptides not in contact with a folded protein domain. We disregarded the disconnected assemblies from the data set. Because authors may indicate asymmetric assemblies that EPPIC may skip, we added the biological assemblies annotated in the PDB to the assembly data set, if they were not present in the EPPIC list for an entry.

The assemblies of a Pfam architectures were clustered based on symmetry, stoichiometry, and the similarity of connecting interfaces (see Methods). So for example, for a given Pfam architecture, D3 hexamers from two different crystals can be compared using two interfaces: one asymmetric interface that defines each C3 cyclic trimer; and one symmetric interface that defines the interaction of two C3 cycles to form a hexamer. A ProtCAD cluster is defined by stoichiometry, symmetry, the number of crystal forms (#CFs) in the cluster. “Crystal forms” are defined as the number of independent experiments for an assembly. Multiple crystallographic entries of the same Pfam have the same crystal form if they have the same space group and similar unit cell dimensions and angles (within 5%). We treat each cryo-EM or NMR structure as a single independent experiment and add them to the #CFs.

If a given protein or protein complex (as defined by a distinct UniProt or UniProts and the Pfam architecture of the construct(s)) is the subject of multiple, independent experiments, it is likely that most of the experiments will contain the same assembly, represented in one of our clusters. There are of course many exceptions due to equilibrium between different assemblies, as well as different experimental conditions, construct definitions, mutations, and ligands. For each UniProt architecture in a cluster, we determined the number of crystal forms in the cluster (CF_UNPclus) and the number of crystal forms for the UniProt and architecture combination across the whole PDB (CF_UNParch). We call the ratio of these two numbers R_CF_UNPclus. For example, the Pfam architecture (Transthyretin) contains 13 UniProts and 27 CFs. The largest D2 tetramer cluster has 12 UniProts and 21 CFs. Since the total number of CFs of these 12 UniProts in the whole PDB is also 21, R_CF_UNPclus is 1.0 (21/21). The second largest D2 tetramer cluster has one UniProt (TTHY_HUMAN) and 5 CFs; TTHY_HUMAN has 7 CFs across the PDB, so the R_CF_UNPclus of TTHY_HUMAN is 0.71 (5/7). R_CF_UNP can be calculated for each UniProt; it can also be calculated for each cluster as the total number of CFs in the cluster with defined UniProts (some chains do not have UniProts) divided by the number of CFs in the whole PDB for those UniProts.

Higher values of R_CF_UNPclus indicate clusters that are more likely to represent biologically relevant assemblies. In many cases, a high R_CF_UNPclus may occur for sub-assemblies of the true assembly (e.g., a homodimer that is part of a D2 homotetramer). The common clusters in ProtCAD with different minimum numbers of CFs and R_CF_UNPclus ≥ 0.7 are summarized in Table 1. For example, there are 1,838 Pfam architectures and 3,291 clusters with at least 5 CFs and R_CF_UNPclus ≥ 0.7, comprising 65,521 PDB entries and 19,955 UniProts. The assemblies in these clusters are represented in the annotated biological assemblies of 85% of the associated entries.

**Table 1.**
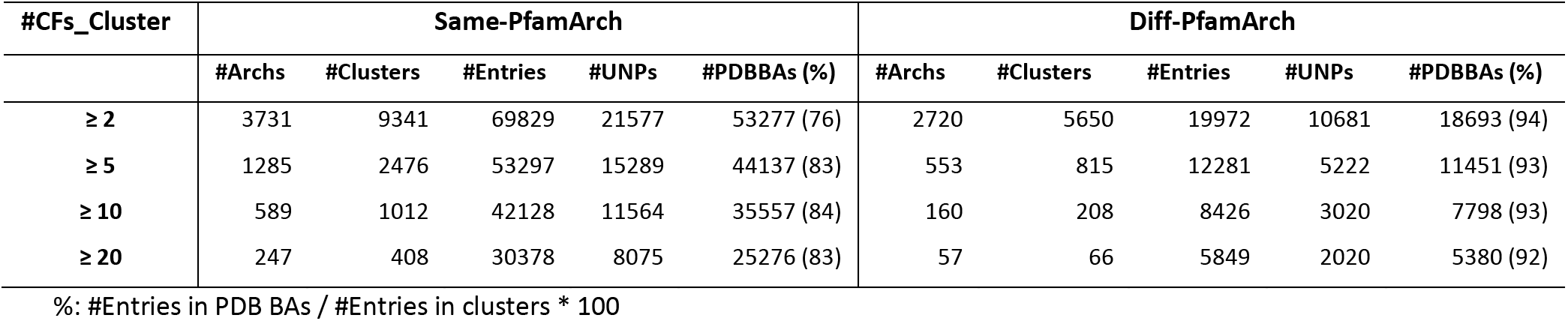
Summary of assembly clusters with R_CF_UNPclus ≥ 0.7

### Symmetry and stoichiometry of common assemblies

The most common form of symmetry in protein assemblies with two or more chains is cyclic symmetry (*Cn*, *n*≥2), which occurs in 51% of all EPPIC assemblies in crystals. This is followed by dihedral symmetry (*Dn n*≥2), which comprises 6% of assemblies. The percentage values are increased to 64% and 15% for *Cn* and *Dn* in common assembly clusters with #CFs_clus ≥ 2 and R_CF_UNPclus ≥ 0.7. Table 2 shows the numbers of Pfam architectures, clusters, entries, and UniProts in cyclic (*Cn*) and dihedral (*Dn*) homomultimers or heteromultimers of a single Pfam architecture, where each cluster has #CFs_clus ≥ 5 and R_CF_UNPclus ≥ 0.7. Each PDB entry is counted only once. If it has several assemblies in Pfam architecture clusters, only the symmetry and stoichiometry of the largest assembly of an entry is used.

**Table 2.** Summary data of cyclic and dihedral *An* assemblies in common clusters (#CFs_clus ≥ 5).

| Symmetry | Stoichiometry | #PfamArchs | #Clusters | #Entries | #Uniprot |
| --- | --- | --- | --- | --- | --- |
| C1 | A2 | 20 | 20 | 1570 | 301 |
|  | A3 | 2 | 2 | 57 | 31 |
|  | A5 | 1 | 1 | 8 | 6 |
|  | A6 | 1 | 1 | 5 | 1 |
| C2 | A2 | 956 | 1222 | 31739 | 9430 |
|  | A4 | 13 | 14 | 841 | 173 |
|  | A12 | 1 | 1 | 5 | 2 |
| C3 | A3 | 125 | 126 | 3425 | 1065 |
|  | A6 | 3 | 3 | 318 | 134 |
|  | A12 | 1 | 1 | 15 | 17 |
| C4 | A4 | 41 | 41 | 1023 | 244 |
|  | A8 | 1 | 1 | 17 | 7 |
| C5 | A5 | 22 | 22 | 616 | 121 |
| C6 | A6 | 15 | 15 | 274 | 82 |
| C7 | A7 | 8 | 8 | 88 | 31 |
| C8 | A8 | 3 | 3 | 19 | 4 |
| C11 | A11 | 1 | 1 | 12 | 4 |
| C12 | A12 | 3 | 3 | 21 | 7 |
| C15 | A15 | 1 | 1 | 6 | 5 |
| C16 | A16 | 1 | 1 | 6 | 2 |
| **Cn** |  | **1219** | **1487** | **40065** | **11667** |
| D2 | A4 | 171 | 184 | 7443 | 2241 |
| D3 | A6 | 58 | 61 | 1618 | 596 |
|  | A12 | 2 | 2 | 132 | 81 |
| D4 | A8 | 21 | 22 | 636 | 233 |
| D5 | A10 | 11 | 11 | 269 | 91 |
| D6 | A12 | 10 | 10 | 138 | 56 |
| D7 | A14 | 3 | 3 | 98 | 40 |
|  | A28 | 1 | 1 | 23 | 8 |
| D8 | A16 | 1 | 1 | 33 | 6 |
| D9 | A18 | 1 | 1 | 5 | 3 |
| D12 | A24 | 1 | 1 | 9 | 3 |
| **Dn** |  | **280** | **297** | **10404** | **3358** |

\* Note: An assemblies may be heterooligomers, if two different protein sequences have the same Pfam architecture.

### Assembly Evolution

Proteins in a Pfam architecture are likely to form the same assemblies when they are closely related in evolution. We analyzed the sequence identities within clusters and between clusters with at least 3 CFs in same-Pfam architectures (Figure 1). While average sequence identities from intra-cluster and inter-cluster protein pairs overlap, the peak of intra-cluster sequence identities is at 50%, while the peak of inter-cluster sequence identities is at 20%. The intra-cluster protein pairs are more homologous than proteins between clusters.

**Figure 1.**
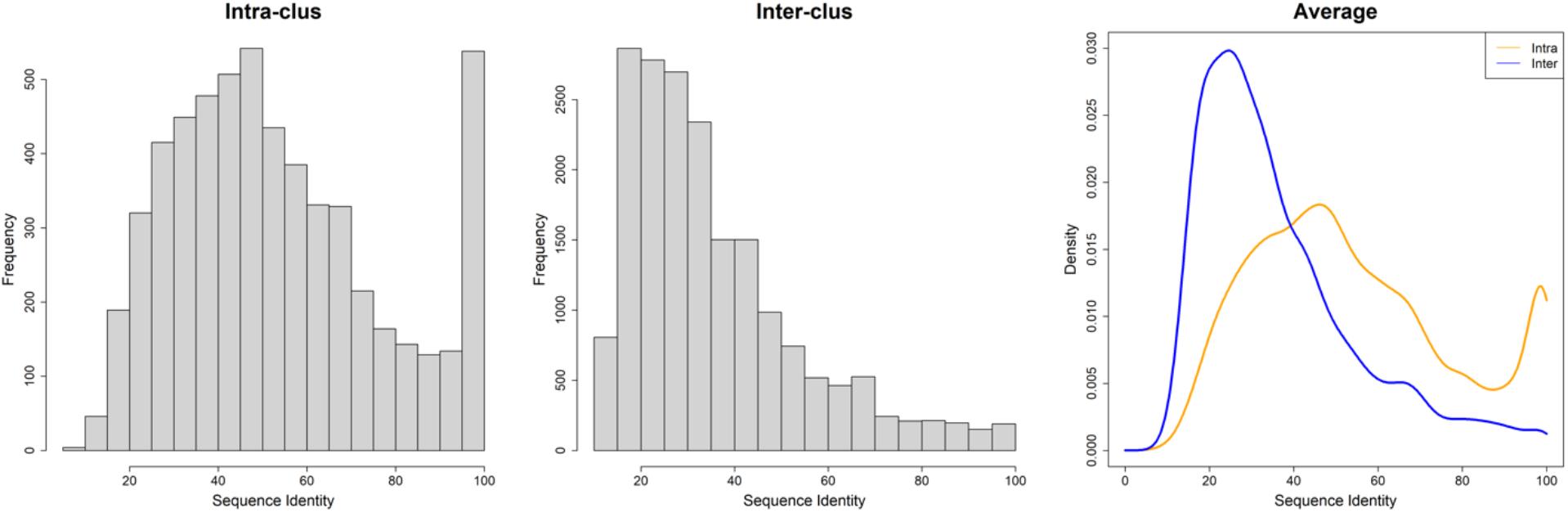
The histogram and density plots of intra-cluster and inter-cluster sequence identities. Clusters are in same-Pfam architectures and contain at least 3 CFs. The Pfam architecture (V-set)_(C1-set) group (antibodies and T-cell receptors) was excluded. Sequence identities are in [0, 100]. The average sequence identities were used in the plots. The left panel is the histogram of intra-cluster average sequence identities, the middle panel is the histogram of inter-cluster average sequence identities, and the right panel is the density functions of intra-cluster (in orange) and inter-cluster (in blue) with bandwidth = 3.

Figure 2 shows an example of Pfam: 4HBT assembly clusters (the enzyme 4HBT synthesizes 4-hydroxybenzoate from 4-chlorobenzoate), which has a common dimer (C2-A2) occurring in 88 CFs, 72 UniProts, and 150 entries. This symmetric dimer also occurs in two different D2 tetramers and one D3 hexamer. The average intra-cluster sequence identity is about 30% while the inter-cluster sequence identity is 17%. The proteins in the two tetramer assembly clusters and the hexamer assembly cluster are colored in green, red, and orange in the phylogenetic tree respectively. The remaining proteins that only have the common dimer assembly are colored in blue. The tetramer in green branches is referred to as the α-tetramer since it is connected by α-helices. The tetramer in red branches is the β-tetramer since it is connected by β-sheets (31). The hexamer is composed of three copies of the common dimer. Two UniProts Q93CG9_9GAMM (PDB: 4I45) and Y1494_AQUAE (PDB: 2EGJ, 2EGR) have both tetramers in their crystals, while the entry 2EGI of Q93CG9_9GAMM and the entry 3R87 of Y1494_AQUAE only have the α-tetramer (green). Evidence from the phylogenetic tree shows that they most likely have the α-tetramer (green). Q2LUI2_SYNAS (PDB: 3HDU) and Q7NVP2_CHRVO (PDB: 4RMM) (no publication for either entry) do not form the β-tetramer (red) in their crystals although they are located in the red branches.

**Figure 2.**
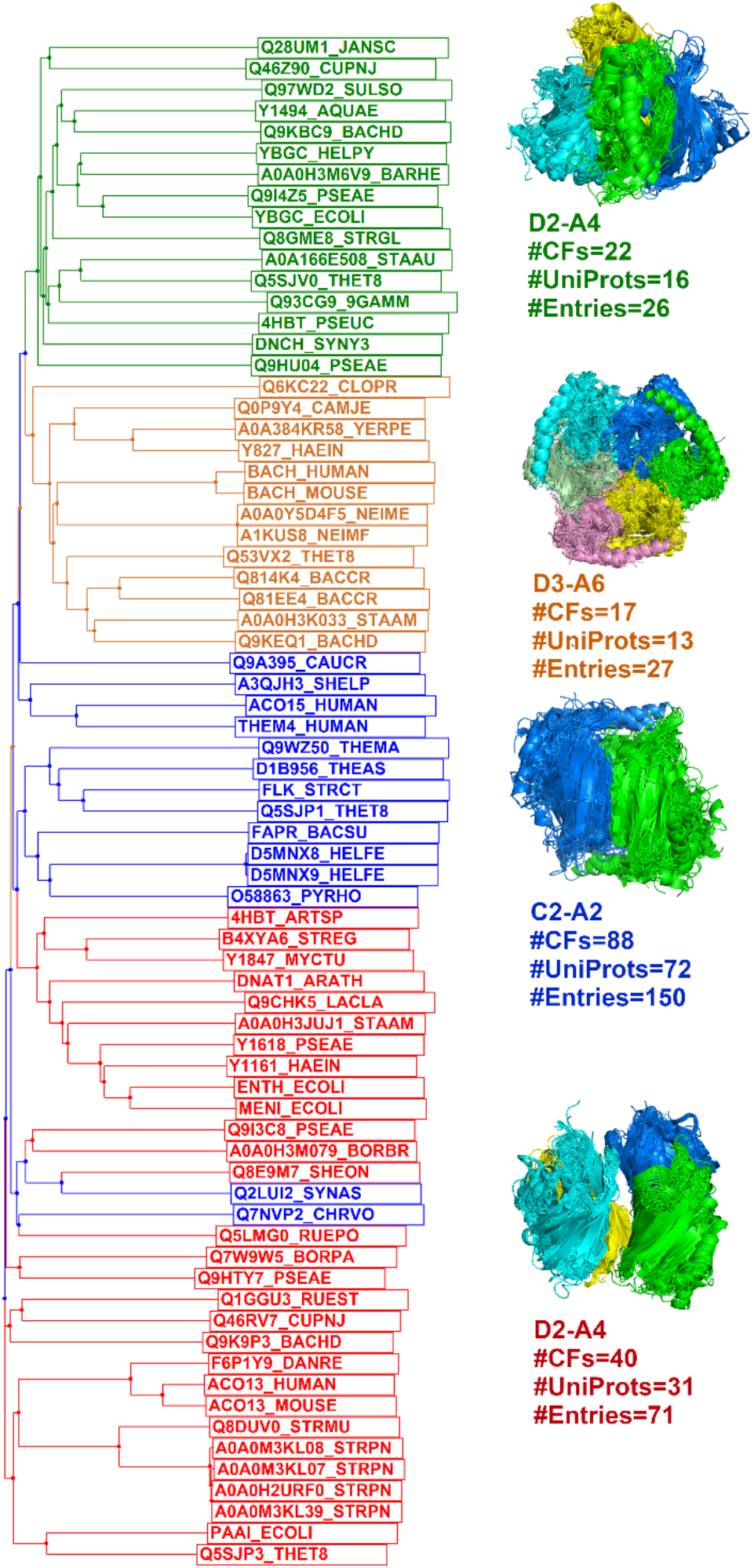
The phylogenetic tree of Pfam 4HBT assemblies. There are two different D2 tetramers (colored in green and red in the tree respectively), one hexamer in D3 symmetry (colored in orange), all of which contain the cluster 1 C2 dimer (side-to-side and head-to-tail, α-helices on one side and β-sheet on the other side, colored in blue) in 88 CFs and 72 UniProts. The tetramer in green branches is formed by α-helical interfaces while the tetramer in red branches is connected by β-sheets. The hexamer is composed of three cluster 1 dimers. The sequence files for UniProt and assemblies can be downloaded from ProtCAD website (http://dunbrack2.fccc.edu/ProtCAD/Results/PfamArchClusterInfo.aspx?GroupId=2822).

### ProtCAD website

The user interface of the ProtCAD web site is presented in Figure 3. There are three types of queries: 1) a PDB ID; 2) a Pfam ID or accession code (e.g., Pkinase or PF00069; and 3) UniProt ID or accession code (e.g., P15056 or BRAF_HUMAN). A PDB ID query will return a table containing the assemblies of this PDB entry, the Pfam architecture, and its clusters. The Pfam architecture is clickable, which leads to an assembly page of this Pfam architecture. Clicking the link “Assembly Clusters” at the top of the page leads to the cluster page. Input of a Pfam ID or a UniProt ID leads to a summary page of Pfam architectures and clusters which contain the input Pfam or Pfams that the UniProt sequence contains. The web table data can be export into a csv file. Clicking a Pfam architecture leads to the cluster page. For UniProt ID input, a web page lists all assemblies containing this UniProt protein in different symmetries and stoichiometries; the coordinates of all the assemblies can be downloaded with a link at the top of the page. A “Browse” page contains all Pfams, Pfam architectures, and UniProts in the PDB with links. Clicking a Pfam ID or a UniProt ID is the same as inputting a Pfam ID or UniProt ID. Clicking a Pfam architecture returns all assemblies in the Pfam architecture. All search and browse pages will lead to the cluster pages. The cluster page is the most functional web page, providing data including clusters, assemblies, and links to coordinates, as well as downloading, sorting, and visualization.

**Figure 3.**
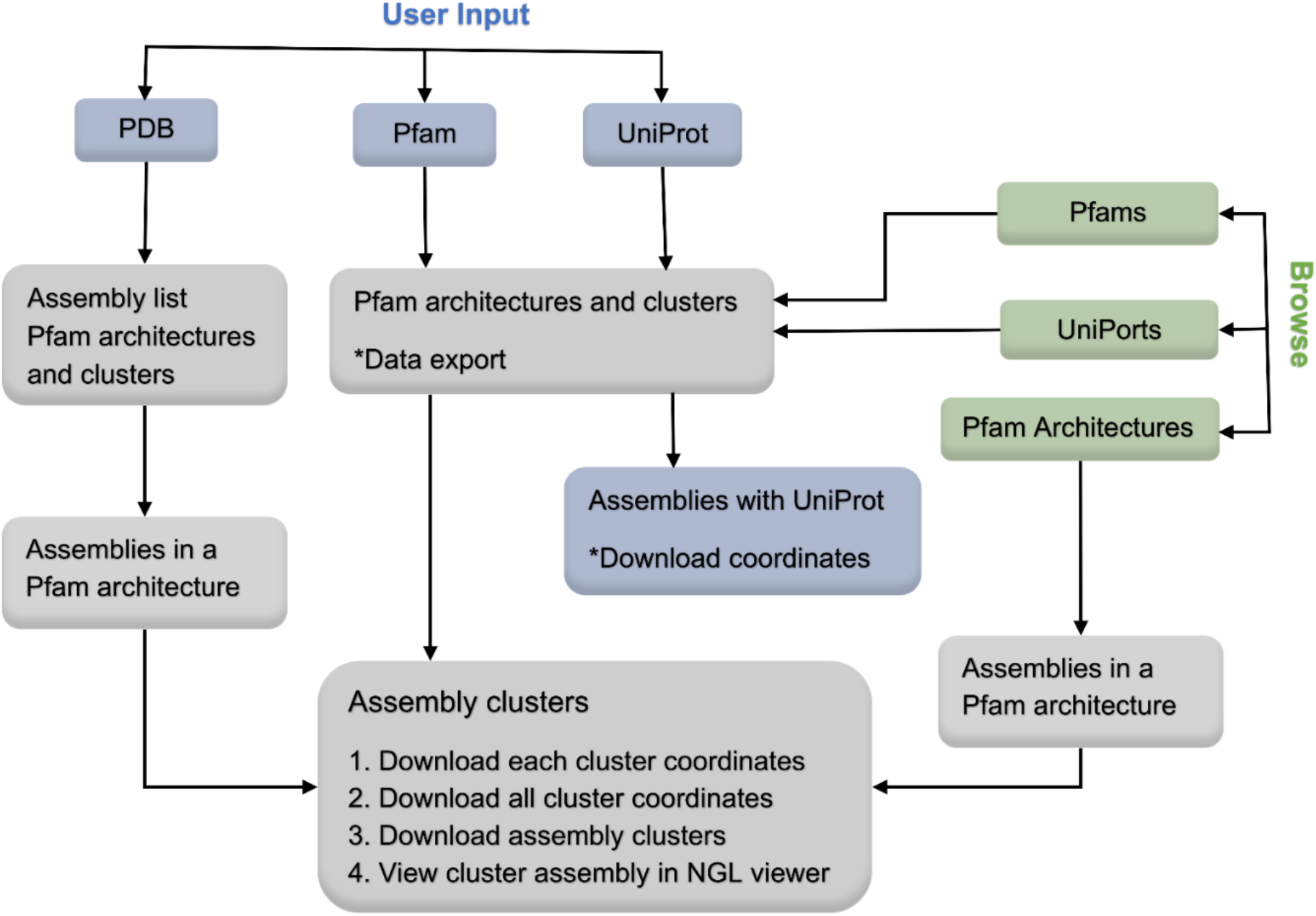
The user interface of ProtCAD web site. A user can input a PDB ID, a Pfam ID, or a UniProt identifier to search assembly clusters to query the ProtCAD database, or browse lists of Pfams, UniProts, and Pfam architectures in ProtCAD. ProtCAD returns a summary table of assembly clusters of one or more Pfam architectures, which leads to the cluster page. Data export, download and visualization are provided in several web pages for the user’s further investigation. The description of each web page and table columns is provided in the web page or the “Help” section of the ProtCAD web site (http://dunbrack2.fccc.edu/ProtCAD/Help/Help.aspx).

As an example, inputting or browsing for the Pfam (IMPDH) or browsing for the Pfam architecture (IMPDH[1-82])_(CBS)_(CBS)_(IMPDH[83-345]) lead to the cluster page of (IMPDH)_(CBS)_(CBS). The assembly cluster page for Pfam architecture (IMPDH)_(CBS)_(CBS) is shown in Figure 4. The (IMPDH) domain is split by two (CBS) domains. The Pfam architecture contains 32 CFs, 11 UniProts, and 47 PDB entries. The user can expand and collapse the assembly table for each cluster, download coordinates and PyMOL scripts of each cluster, view images of clusters by mousing over the thumbnail images, or view a cluster assembly in NGL viewer by clicking a thumbnail image. The cluster table can be sorted by selecting a header in the dropdown box at the middle top of the page. The data of assembly clusters in the text file (e.g. (IMPDH[1-82])_(CBS)_(CBS)_(IMPDH[83-345])_349.txt.gz) and a tar file containing coordinates for all assemblies in the cluster (e.g. (IMPDH[1-82])_(CBS)_(CBS)_(IMPDH[83-345])_349.tar) can also be downloaded from the links at the top right corner of the page. The (IMPDH)_(CBS)_(CBS) cluster page shows 18 clusters with each containing at least 2 entries. Although a common cluster must contain at least 2 CFs, a cluster page displays clusters with 1 CF and ≥ 2 entries so that some clusters with large assemblies can also be presented, like *A24* in octahedral symmetry (O) in this page. Generally, there are fewer large assemblies, and they might be interesting to know and are likely to form small clusters.

**Figure 4.**
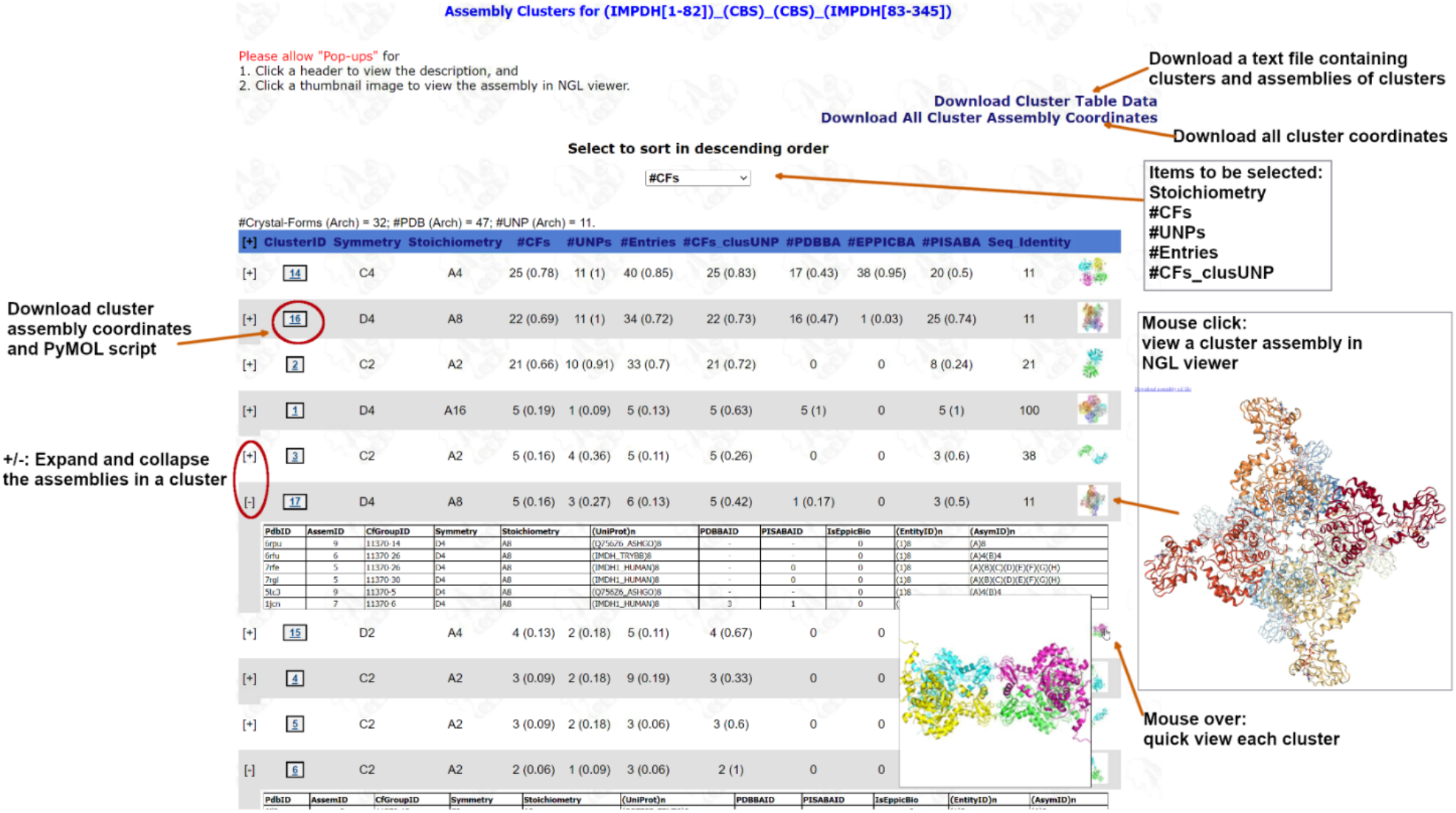
Screenshot of the cluster web page for Pfam architecture IMPDH_CBS_CBS. The IMPDH domain is split by two CBS domains. The numbers in parentheses are the ratio between the number of CFs, UniProts, and Entries in a cluster and the number each of these in the whole PDB for the same Pfam architecture. UNPclus means the number of CFs of UniProts in a cluster (which may be less than the number of CFs in the cluster, because not all chains and CFs have UniProts in the PDB), the ratio in parentheses is therefore the #CFs_UNPclus divided by #CFs_UNParch (the number of CFs for those UniProts/Pfam architecture across the whole PDB). For #PDBBA, #EPPICBA and #PISABA, the ratio is #entries where the cluster assembly is same as PDB/EPPIC/PISA biological assemblies divided by #Entries of the cluster. To view description about a header or view a cluster assembly in NGL viewer (https://github.com/nglviewer/nglview), “Pop-ups” must be allowed in the web browser.

### Examples

#### Same-Pfam architecture

Inosine-5’-monophosphate dehydrogenase (IMPDH) is a key regulatory enzyme in purine nucleotide biosynthesis, has been recognized as a target for cancer and viral chemotherapy since its activity is tightly linked with cell proliferation (32,33). IMPDH can assemble filaments in response to changes in metabolic demand (34). Pfam architecture (IMPDH)_(CBS)_(CBS), described above, has two D4 octamers with ≥ 5 CFs (Figure 5A and 5B) which are composed of the (IMPDH) filament (Figure 5C). The largest D4 octamer occurs in 23 CFs, 11 UniProts, and 35 entries (Figure 5A), including both active/extended and inactive/compressed conformations. There are five octamers (IMDH1_HUMAN: 1JCN, 7RES, 7RFH and 7RGM; Q9HSM5_PSEAE: 4DQW,) in an active conformation. The second D4 octamer cluster comprises 5 CFs, 3 UniProts, and 6 entries (Figure 5B); among them, there are 2 EM structures (PDB: 7RFE and 7RGL) of human IMPDH1. Four out of six entries (PDB: 1JCN, 5TC3, 6RFU and 6RPU) are crystals containing both octamers, which can be used to build a model of an IMPDH filament. The filament was built from PDB: 6RFU by overlapping two D4 octamers. The filament model is in an inactive/compressed conformation. The largest cluster is a C4 tetramer (Figure 5D) which is the common sub-structure in the two D4 octamers by face-to-face or back-to-back interactions, the D4 hexadecamer (A16) (Figure 5E), and the octahedral A24 assembly (Figure 5F). The D4 hexadecamer contains 8 full-length IMPDH2 chains and 8 peptides (colored in magenta) which are fragments of the full-length IMPDH2 chains, which is all that is observed in the cryo-EM electron density of the filament; the global symmetry is D4 based on the full protein octamer, which is same as the cluster in Figure 5A. IMDH_TRIFO forms a unique octahedral 24-mer that exists in 7 entries, with two CBS domains are disordered.

**Figure 5.**
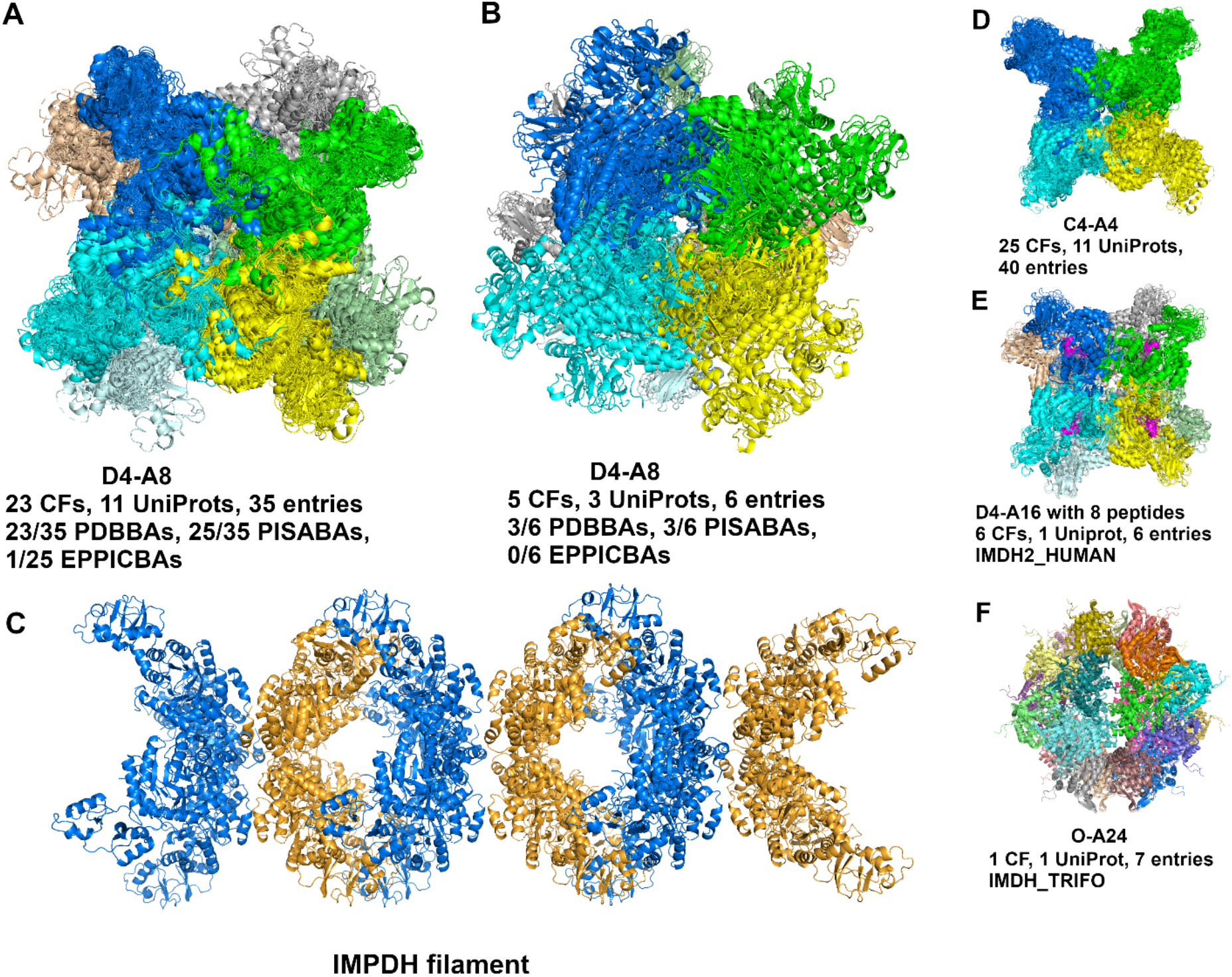
Examples of same-Pfam architecture IMPDH_CBS_CBS clusters. (**A**) The largest D4-A8 cluster occurs in 23 crystal forms, 11 UniProts, and 35 PDB entries. This D4 octamer is in 23 PDB biological assemblies, 25 PISA biological assemblies, and one that is marked as “BIO” in EPPIC. There are 5 octamers in the extended/active conformation, including 4 entries (PDB: 1JCN, 7RES, 7RFH and 7RGM) of IMDH1_HUMAN and 1 entry (PDB: 4DQW) of Q9HXM5_PSEAE. Except 1JCN, each is bound with 16 ATPs (one on each CBS domain). (**B**) The second D4-A8 cluster occurs in 5 crystal forms, 3 UniProts, and 6 PDB entries. The cluster contains two EM structures: 7RFE and 7RGL of IMDH1_HUMAN, and four crystals of PDB: 1JCN (IMDH1_HUMAN), 5TC3 (Q756Z6_ASHGO), 6RFU (IMDH_TRYBB) and 6RPU (Q756Z6_ASHGO). (**C**) The structure of IMPDH filament contains two D4-A8 clusters, built from PDB: 6RFU. (**D**) the C4-A4 assembly is the sub-structure of clusters in (A), (B), (E) and (F). The first D4 octamer in (A) contains two face-to-face C4 tetramers. The second D4 octamer contains two back-to-back C4 tetramers. (**E**) D4-A16 cluster contains 8 copies of IMPDH peptides, the super-stoichiometry is A8, which is the same as D4-A8 in (A). (**F**) The octahedral-symmetry A24 cluster exists in one crystal form of IMDH_TRIFO and 7 PDB entries.

#### Diff-Pfam architecture

ProtCAD contains 2853 diff-Pfam architectures with clusters with at least 2 CFs. A user can input a Pfam ID or select a Pfam ID in the “Browse” page, to view all Pfam architectures containing the Pfam ID, or a user can directly select a Pfam architecture in the “Browse” pages, then click to go to the cluster page about this Pfam architecture. As an example, if a user’s input is the Pfam (FGF) and the Pfam architecture (FGF)(I-set)_(I-set) is selected, it leads to the cluster page. The Pfam (I-set) domains comprise the Ig-like II and Ig-like III domains of FGFRs. FGF/FGFR signaling plays essential roles in development, metabolism, and tissue homeostasis. Malfunction of this signaling pathway leads to various diseases including cancers (35). FGF and FGFR extracellular regions are known to form a minimal complex with 2:2 stoichiometry (36) but are also able to form larger species in the presence of heparan sulfate (37). Figure 6 shows a 2:2 heterotetramer with C2 symmetry that contains two FGF chains and two FGFR chains (Figure 6A). The cluster contains the entries: FGF10/FGFR2 (PDB:1NUN); FGF1/FGFR2 (4J23, 1E0O); FGF2/FGFR1 (1CVS, 1FQ9) in 5 CFs. PDB entry 1E0O has different conformation of its Ig-like III domains, as shown in Figure 6B. The known C2 heterotetramer in Figure 6A occurs in the PDB biological assemblies of two entries (FGF2/FGFR1 in 1FQ9, 1CVS) but is missing in the FGF10/FGFR2 and FGF1/FGFR2 entries (1NUN, 4J23, 1E0O). The only biological assembly in the PDB for 1E0O is the same as the asymmetric unit (Figure 6B inset), but is different from the assembly in the cluster which more closely resembles other FGF/FGFR heterotetramers. The same set of entries contains a cluster of octameric (4:4) assemblies with D2 symmetry with interactions between the II and III domains of FGFR of one tetramer with another. The biological relevance of this assembly is not known.

**Figure 6.**
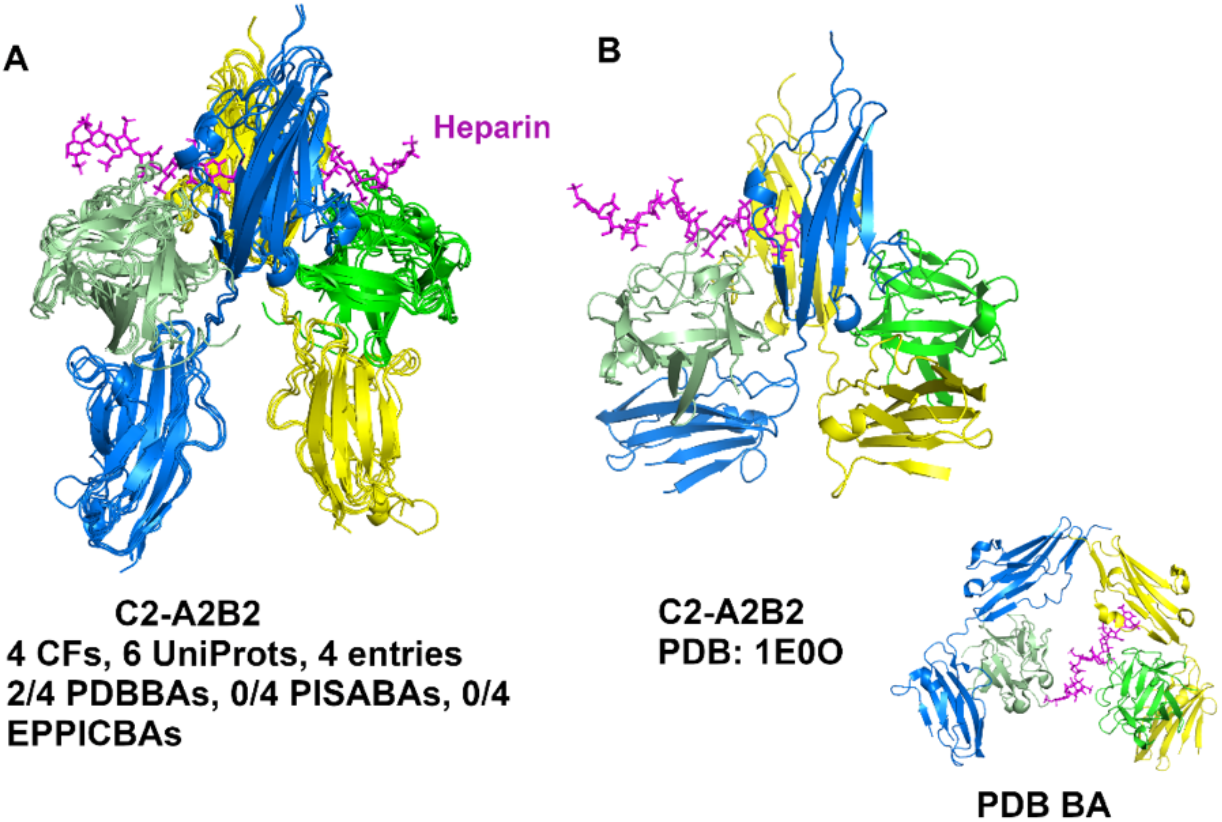
Two clusters of assemblies of FGF and Ig domains of FGF receptor (FGFR). Heparin is colored in magenta. The FGF chains are colored in green and light green in all panels. The FGFR Ig chains are colored in blue and yellow in (A) and (B). Ig chains are colored in blue, yellow, orange, and light blue in (C) and (D). (A) C2 tetramer cluster without PDB: 1E0O. (B) The C2 tetramer of PDB: 1E0O. The inset is the PDB biological assembly generated by the PQS program, and is the same as the asymmetric unit (https://www.rcsb.org/structure/1E0O). (C) D2 octamer cluster without PDB: 1E0O. (D) The D2 octamer of PDB: 1E0O.Figure 6 show C2 heterotetrameric assembly cluster of FGF and FGFR Ig-like II and Ig-like III domains.

#### UniProt Search

A user can input or browse a UniProt ID to search ProtCAD. As an example, Figure 7 presents a Pkinase dimer cluster with 20 CFs, 13 UniProts and 51 entries containing 8 entries of WEE1_HUMAN. WEE1_HUMAN is a nuclear Ser/Thr kinase (Pfam: Pkinase) and is a key regulator of cell cycle progression. The N-terminal dimer of mouse PERK (E2AK3_MOUSE, PDB: 3QD2) was reported to perform autophosphorylation by inter-dimer interactions, and this mechanism may be shared amongst all eIF2α family members (38). The cluster shows that it occurs in all eIF2α family entries in the PDB, including PKR (E2AK2_HUMAN: 2CFs, 3 entries), PERK (E2AK3_HUMAN: 4 CFs, 10 entries; E2AK3_MOUSE: 1 CF, 1 entry) and GCN2 (E2AK4_HUMAN: 2 CFs, 3 entries). R_CF_UNPclus is therefore equal to 1.0 for these proteins. The average similarity Q score of the dimer of WEE1_HUMAN with the dimers of PKR, PERK and GCN2 is 0.68, a highly confident value, while the average sequence identity is 24%. The RMSD is 3.618 Å over 471 residues when superposing the WEE1 dimer on the E2AK3 dimer. In our phylogenetic tree for human protein kinases based on a structurally validated multiple sequence alignments (39), WEE1 occurs on the same branch as PKR, PERK, and GCN2.

**Figure 7.**
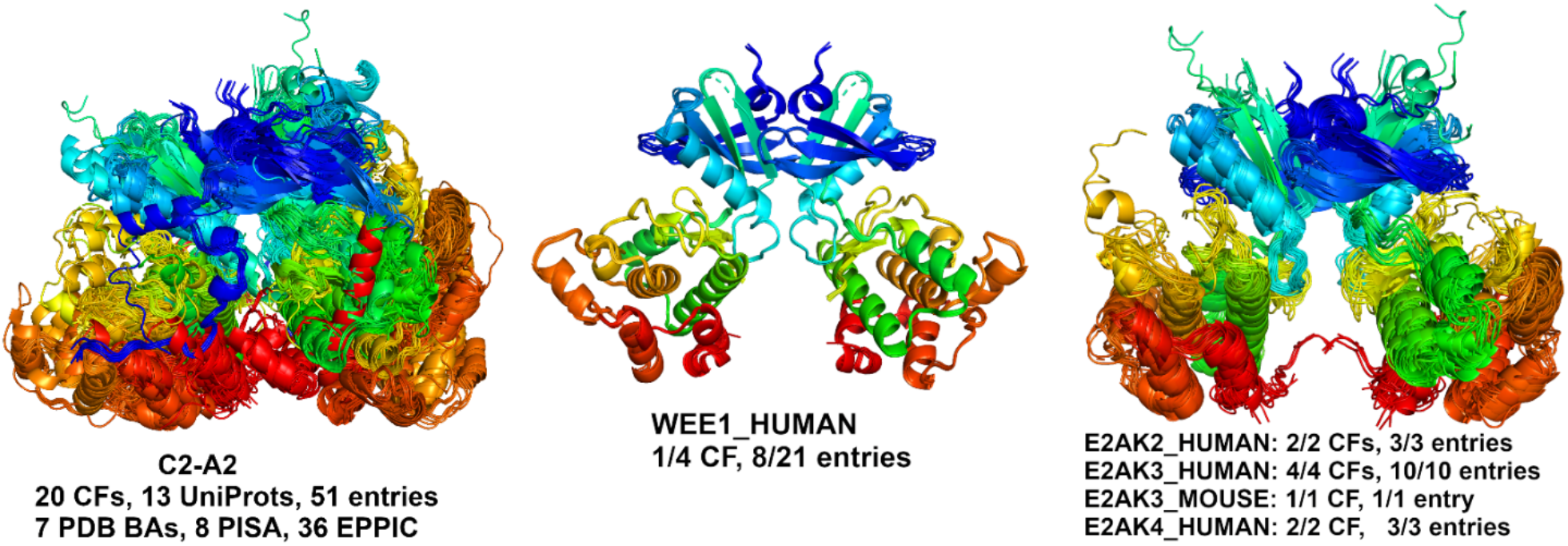
Pkinase dimer cluster containing a symmetric dimer of WEE1_HUMAN. A MAPK2_HUMAN dimer was removed due to poor superposition and is likely a chance similarity. The dimer occurs in all crystal forms and PDB entries of E2AK2_HUMAN (PKR), E2AK3_HUMAN (PERK), E2AK3_MOUSE (PERK) and E2AK4_HUMAN (GCN2).

## DISCUSSION

One of the unsolved problems in structural biology is how to determine the correct biological assembly for proteins in crystallographic structures. Previous studies show the error rates are significant in three major biological assembly sources: PDB, PISA, and EPPIC. It is estimated that up to 17% of the X-ray structures in the PDB have one or more incorrect annotations of the most likely biological assembly (9). The error rates for PISA and EPPIC, two major biological assembly sources, are even higher: 13% and 18% for homodimers and 16% and 32% for larger homooligomeric complexes (6), 25% and 33% for heterodimers, and 39% and 45% for larger hetero complexes respectively (40). The inconsistency rates for PDB, PISA, and EPPIC from our ProtCAD common clusters with at least 5 CFs are 15%, 23% and 24% respectively. This inconsistency has been widely acknowledged to be a very important issue that affects structural biological analysis and computational prediction on protein complexes. We provide ProtCAD as an alternative and complementary source of information on the biological assemblies in crystal structures. When there is some doubt about an assembly, e.g., inconsistency among different sources, observing the same assembly across independent experiments of the same or homologous proteins may provide evolutionary and physical evidence in favor of that assembly. One important function of ProtCAD is to display representative structures of the assembly clusters of homologous proteins on each Pfam architecture group web page. Thus it is straightforward to view various possible assemblies and the independent experimental sources of those assemblies (different crystal forms and cryo-EM and NMR experiments).

Since we build assemblies with the EPPIC program, rather than taking the biological assemblies deposited in the PDB, our data can provide information on previously unrecognized biological assemblies. For example, we observe a common dimer in the closely related human and mouse kinases WEE1, PKR, PERK, and GCN2 (the last three are EIF2AK2, EIF2AK3, EIF2AK4) (Figure 7). One paper on mouse PERK demonstrates the biological relevance of the dimer (38), while the rest ignore it. WEE1, which regulates entry into mitosis, is an important therapeutic target in cancer (41). We also showed that some well-known assemblies, such as the 2:2 complex of FGF and FGFR extracellular domains, are present in some entries but not present in any of the PDB’s biological assemblies or even shown in the relevant publications. ProtCAD can be also used to combine assemblies that may lead to the identification of larger assemblies or filaments. The example of IMPDH shows that the two different D4 octamers can be combined to form an IMPDH filament (Figure 5).

A limitation of ProtCAD is that we do not provide probabilities that the assemblies in any particular cluster are the correct biological assemblies. This is for two reasons. First, there is simply not enough benchmark data of “true” biological assemblies for protein crystal structures that is not derived from methods related to those of ProtCID and ProtCAD. To use ProtCID data might involve circular reasoning if it is not done carefully enough. Second, some assemblies may be transient but are observable at the high concentrations needed for X-ray crystallography. But such assemblies might not be scored with high probability and a potentially valid hypothesis might be neglected. It is possible that deep learning predictors may reproduce these assemblies, providing two forms of evidence in their favor. Using AlphaFold-Multimer (42) on ColabFold (43), for example, we were able to reproduce almost exactly the dimers of WEE1, PKR, PERK, and GCN2 kinases shown in Figure 7.

ProtCAD is the first comprehensive structural database comparing and clustering potential biological assemblies across all publicly available experimental structures of homologous proteins. Recently, our ProtCID database has been widely used for benchmarking interface/assembly predictors in the 3D-BioInfo community of ELIXIR (https://elixir-europe.org/communities/3d-bioinfo). Similarly, we believe the common assembly clusters in ProtCAD can also be used for benchmark data, as well as training and testing data sets for structure prediction of protein complexes, especially in the rapidly developing field of deep learning structure predictors (42,44,45).

## DATA AVAILABILITY

The ProtCAD webserver is available at http://dunbrack2.fccc.edu/protcad.

